# Two doses of mRNA vaccine elicit cross-neutralizing memory B-cells against SARS-CoV-2 Omicron variant

**DOI:** 10.1101/2021.12.24.474091

**Authors:** Ryutaro Kotaki, Yu Adachi, Saya Moriyama, Taishi Onodera, Shuetsu Fukushi, Takaki Nagakura, Keisuke Tonouchi, Kazutaka Terahara, Lin Sun, Tomohiro Takano, Ayae Nishiyama, Masaharu Shinkai, Kunihiro Oba, Fukumi Nakamura-Uchiyama, Hidefumi Shimizu, Tadaki Suzuki, Takayuki Matsumura, Masanori Isogawa, Yoshimasa Takahashi

## Abstract

SARS-CoV-2 Beta and Omicron variants have multiple mutations in the receptor-binding domain (RBD) allowing antibody evasion. Despite the resistance to circulating antibodies in those who received two doses of mRNA vaccine, the third dose prominently recalls cross-neutralizing antibodies with expanded breadth to these variants. Herein, we longitudinally profiled the cellular composition of persistent memory B-cell subsets and their antibody reactivity against these variants following the second vaccine dose. The vaccination elicited a memory B-cell subset with resting phenotype that dominated the other subsets at 4.9 months. Notably, most of the resting memory subset retained the ability to bind the Beta variant, and the memory-derived antibodies cross-neutralized the Beta and Omicron variants at frequencies of 59% and 29%, respectively. The preservation of cross-neutralizing antibody repertoires in the durable memory B-cell subset likely contributes to the prominent recall of cross-neutralizing antibodies following the third dose of the vaccine.

**One Sentence Summary:** Fully vaccinated individuals preserve cross-neutralizing memory B-cells against the SARS-CoV-2 Omicron variant.

## INTRODUCTION

SARS-CoV-2 mRNA vaccination elicits circulating antibodies from plasma cells and memory B (B_mem_) cells, which constitute two layers of the humoral protection against infecting viruses (*1–8*). The pre-existing antibodies provided by plasma cells prevent infection at the initial infection site and contribute to sterilizing immunity if they are present in sufficient amounts at the time of infection. However, the pre-existing antibodies wane with time after vaccination with mRNA vaccines (*1, 2, 9*). The viruses or viral antigens are then captured by B_mem_ cells that robustly supply plasma cells as the second arm of humoral protection. Importantly, B_mem_ cells remain in the body without noticeable decline for up to six months after two vaccine doses (*1, 2*). Therefore, robust humoral protection by persistent B_mem_ cells along with T-cell-mediated cellular immunity is expected to prevent the onset of symptomatic or severe disease in vaccinees (*10*), which is supported by the long-lasting vaccine effectiveness in preventing severe disease (*10*).

The emergence of SARS-CoV-2 variants raises the issue of immune evasion, as some of them carry intensive mutations in the receptor-binding domain (RBD) of the spike protein, the main target for potently neutralizing antibodies. Among the variants that have emerged so far, Beta and Omicron variants have a greater ability to escape from neutralizing antibodies than Alpha and Delta variants, that are dominantly circulated in the past (*11*). In particular, the Omicron variant drastically reduced the neutralizing activity of circulating antibodies in fully vaccinated individuals and many therapeutic monoclonal antibodies in clinical use (*12–16*). Epidemiological evidence further suggest a reduction in the vaccine effectiveness against this variant (*17*).

Despite the antigenic alteration in the variants, the third dose of mRNA vaccines recalls the cross-neutralizing antibodies that are more resistant to Beta and Omicron variants than the antibodies elicited after the second vaccine dose (*16, 18, 19*). One possible explanation for this phenomenon is that the B_mem_ cells induced after the second vaccine dose are more cross-reactive against the variants than the circulating antibodies from plasma cells. Therefore, to understand the immunological basis behind this phenomenon, it is important to dissect the cross-reactive B_mem_ cell response elicited by two doses of SARS-CoV-2 mRNA vaccines, focusing on the breadth against Beta and Omicron variants and the cross-neutralizing activity.

In this study, we observed a temporal shift in the composition of the B_mem_ cell subsets over time in those who have received two doses of mRNA vaccine. One of the persistent B_mem_ cell subsets with resting phenotype durably preserved cross-reactive antibody repertoires up to 4.9 months after vaccination and secreted cross-neutralizing antibodies against both Beta and Omicron variants when stimulated *in vitro*. Thus, despite the significant RBD mutations in emerging variants, mRNA vaccine based on the original Wuhan strain is able to elicit B_mem_ cell subset with breadth against the variants. Therefore, booster vaccination that restimulates such cross-neutralizing B_mem_ cells potentiates the protective immunity against the infection by SARS-CoV-2 variants.

## RESULTS

### Study cohort

We initially recruited 51 healthcare workers who received two doses of the Pfizer-BioNTech BNT162b2 mRNA vaccine for longitudinal blood donation. During the early (median = 31 days) and late (median = 146.5 days) time points after the second dose, the circulating antibodies and B_mem_ cells in the peripheral blood were analyzed using quantitative and qualitative parameters (fig. S1A). Based on the prior history of COVID-19 diagnosis and anti-nucleocapsid antibodies in prevaccinated plasma, 40 volunteers were selected as naïve for further analysis (fig. S1B). The median age was 46.5 years, ranging from 25 to 73 years, and 16/40 volunteers (40%) were male (fig. S1C). No significant difference was observed in the median age between males (*n* = 16, median: 45.5, IQR: 35.25–57.5) and females (*n* = 24, median: 47, IQR: 36.25–51).

### Temporal maturation of plasma neutralizing antibody

In COVID-19 convalescent individuals, we and others have previously revealed that neutralizing antibodies in plasma improve the potency and breadth against the variants over time, enhancing the resilience of neutralizing antibodies against variants (*20, 21*). Herein, we assessed the temporal maturation of vaccine-elicited neutralizing antibodies using multiple serological parameters. We focused on RBD, as it includes the major epitopes of neutralizing antibodies (*22–27*). The concentration of RBD-binding IgG and IgA antibodies in plasma were quantified by electrochemiluminescence immunoassay (ECLIA) and expressed as binding arbitrary units (BAU) per mL. The neutralizing antibody titers were determined by neutralization assay using authentic viruses and expressed as the endpoint titers that blocked the virus cytopathic effects (*20*). For the initial set of experiments, RBD proteins were prepared from the vaccine strain (Wuhan), Beta variant, and Delta variant with less antigenic changes. Consistent with previous reports, two doses of mRNA vaccine robustly induced Wuhan RBD IgG titers, increasing at one month > 2000-folds above the level of pre-vaccination (Fig. 1A and fig. S2A). Although the Wuhan RBD IgA titers in plasma increased 44-folds after vaccination (fig. S2A), the IgG dominance and better correlation with neutralizing antibody titers suggest that IgG antibodies are major contributors to the neutralizing activity of plasma antibodies (fig. S2B).

**Fig. 1.**
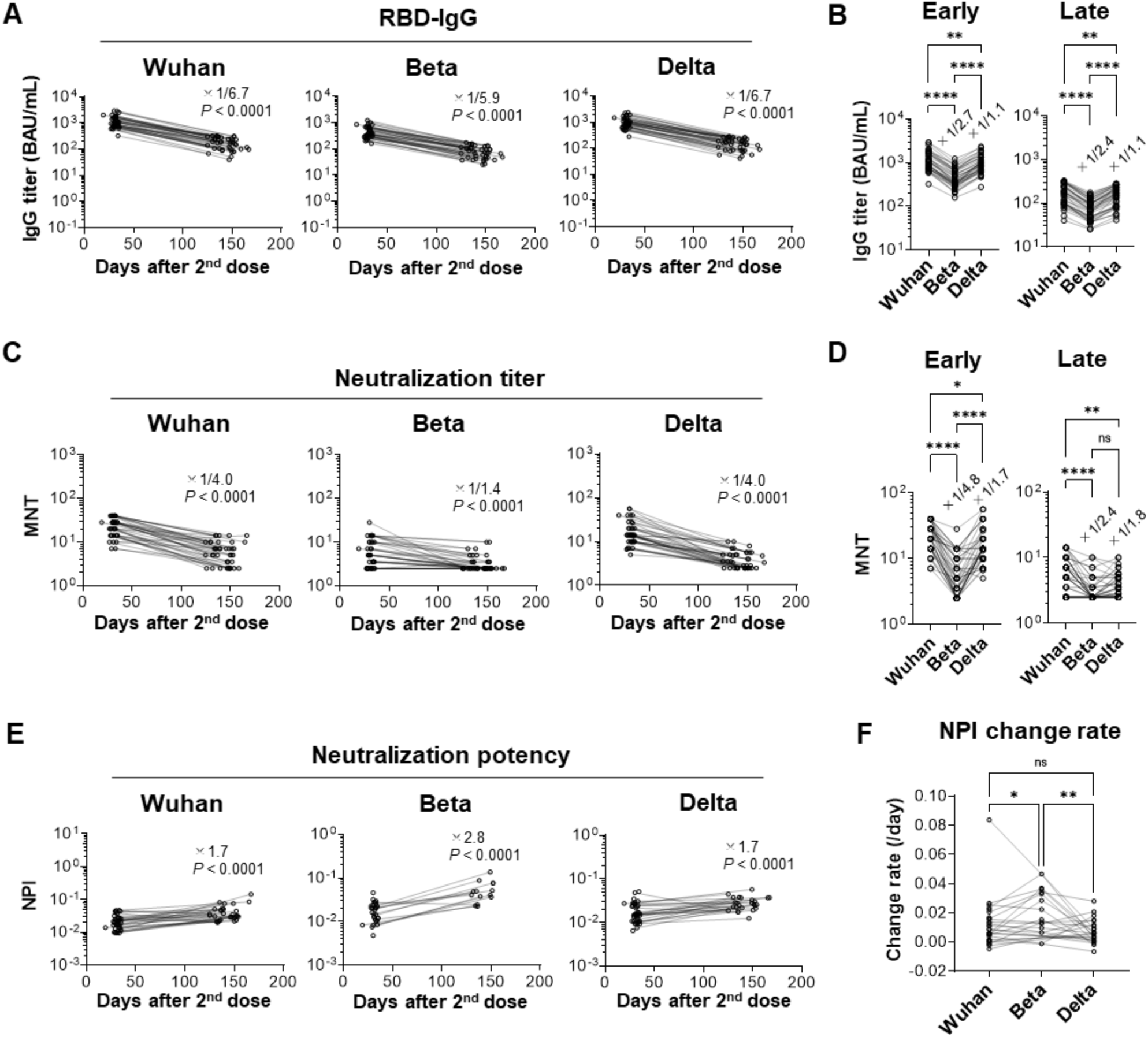
Longitudinal analysis of neutralizing potency of plasma antibody after mRNA vaccination. **(A)** RBD-binding IgG titers in plasma were quantified by ECLIA and expressed in binding arbitrary unit (BAU)/mL. (**B**) The fold-changes of RBD IgG titers were calculated as the titers against the variant divided by the titers against the Wuhan strain. **(C)** Neutralizing antibody titers (MNT) were quantified using authentic viruses. (**D**) The MNT fold-changes were calculated as the titers against the variant divided by that against Wuhan strain. **(E)** Neutralizing potency indices (NPIs) were calculated by dividing MNT by RBD IgG titers from individual virus variant. Samples with the neutralizing titers below the detection limit (2.5) were excluded from the analyses. (**F**) NPI changing rate per day from the early to the late time point were plotted. Median of the fold-changes (A and C) or fold-change of median (E) are indicated with *P* values analyzed with the Wilcoxon test (A and C) or Mann-Whitney U-test (E). The Friedman test was performed in (B and D). For the changing rates (F), statistical analysis was performed with the Kruskal-Wallis test (\**P* < 0.05, \*\**P* < 0.01, \*\*\**P* < 0.001, \*\*\*\**P* < 0.0001, n.s.: not significant).

Vaccine-elicited IgG antibodies also bound to Beta and Delta RBD, albeit at a significant reduction (2.4-2.7-folds for Beta and 1.1-fold for Delta) (Fig. 1B). Neutralizing antibodies against authentic viruses were more severely attenuated by variant mutations (Fig. 1, C and D). Similar to the results obtained with convalescent plasma (*20*), the neutralizing antibodies against the Beta variant declined more slowly than those against Wuhan and Delta (Fig. 1, C and D). Longer persistence of the neutralizing antibodies than that of RBD IgG resulted in a temporal increase in the neutralization potency index (neutralization potency per RBD IgG) with a higher increase in Beta than in Wuhan and Delta strains (Fig. 1, E and F). Thus, a gradual and prominent increase in neutralizing potency over time was observed against the Beta variant in mRNA vaccinees, resembling that of convalescent individuals.

The breadth of circulating antibodies against the Beta and Delta variants was evaluated based on RBD-binding IgG titers. The IgG breadth was calculated by dividing the Beta or Delta RBD IgG by the Wuhan RBD IgG. We observed a 1.1-fold increase in the IgG breadth against the Beta variant from the early to the late time points (Fig. 2A). The breadth of neutralizing antibodies, denoted as neutralizing breadth index (NBI), increased two-fold against Beta variant but not against Delta (1.1-fold decrease) variant (Fig. 2B). A similar increase in NBI was reproduced by pseudovirus (PV) neutralization assay in which neutralization antibody titers were assessed as the half-maximal inhibitory concentrations (IC50) (Fig. 2C and fig. S3). In summary, based on the multiple parameters of serological analysis, the neutralizing potency and breadth against Beta but not Delta variants gradually increased following mRNA vaccination, suggesting the qualitative maturation of circulating antibodies as previously observed in convalescent plasma (*20*).

**Fig. 2.**
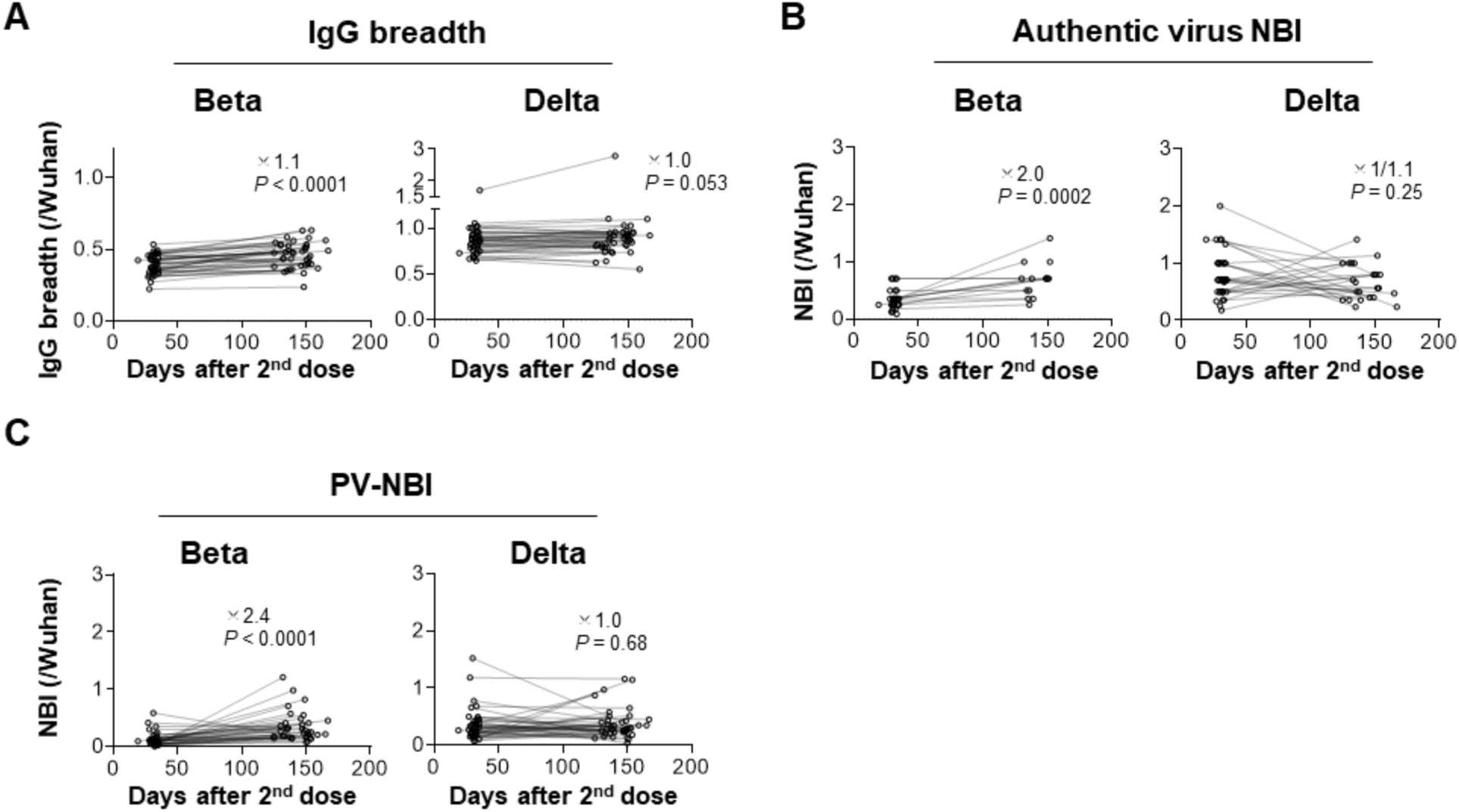
Longitudinal analysis of neutralization breadth of plasma antibody after mRNA vaccination. **(A)** Breadth of RBD IgG calculated as the titer against the variant divided by that against the Wuhan strain were plotted. (B) Breadth of MNT calculated as the titer against the variant divided by that against the Wuhan strain were plotted as neutralization breadth index (NBI). Samples with the neutralizing titers below the detection limit (2.5) at either the early or the late time points were excluded from the neutralization breadth analysis. (C) Breadth of neutralizing antibody titers detected by VSV-pseudoviruses (PV) were calculated similarly as described in (A) and plotted as NBI. Median of the fold-changes (A) or fold-change of median (B) are indicated with P values analyzed with the Wilcoxon test (A) or Mann-Whitney U-test (B).

### Temporal shift of B_mem_ cell subsets in vaccinees

We then tracked the frequencies and phenotypes of RBD-binding B_mem_ cells in peripheral blood via flow cytometry. RBD-binding B-cells in all samples were determined by simultaneous labeling with the Wuhan spike and RBD probes that were coupled to different fluorescent dyes. After gating on spike/RBD-binding CD19^+^CD20^+^IgM^-^IgG^+^ B_mem_ cells, the cells were further subdivided into CD27^+^CD21^+^ (resting), CD27^low/-^CD21^+^ (CD27^low^), CD27^+^CD21^-^ (activated), and CD27^-^CD21^low^CD11c^+^FcRL5^+^ (atypical) (fig. S4) (*28, 29*). Despite the quantitative decline in circulating IgG antibodies in the vaccinees, the number of RBD-binding IgG^+^ B_mem_ cells increased at 1.8-fold from the early to late time points (Fig. 3A), corresponding with previous reports (*1, 2*). In contrast, IgA^+^ and IgM^+^ B_mem_ cells were less persistent than IgG^+^ B cells. Hereafter, we focused on RBD-binding IgG^+^ B_mem_ cells that were robustly induced and durably maintained after mRNA vaccination.

**Fig. 3.**
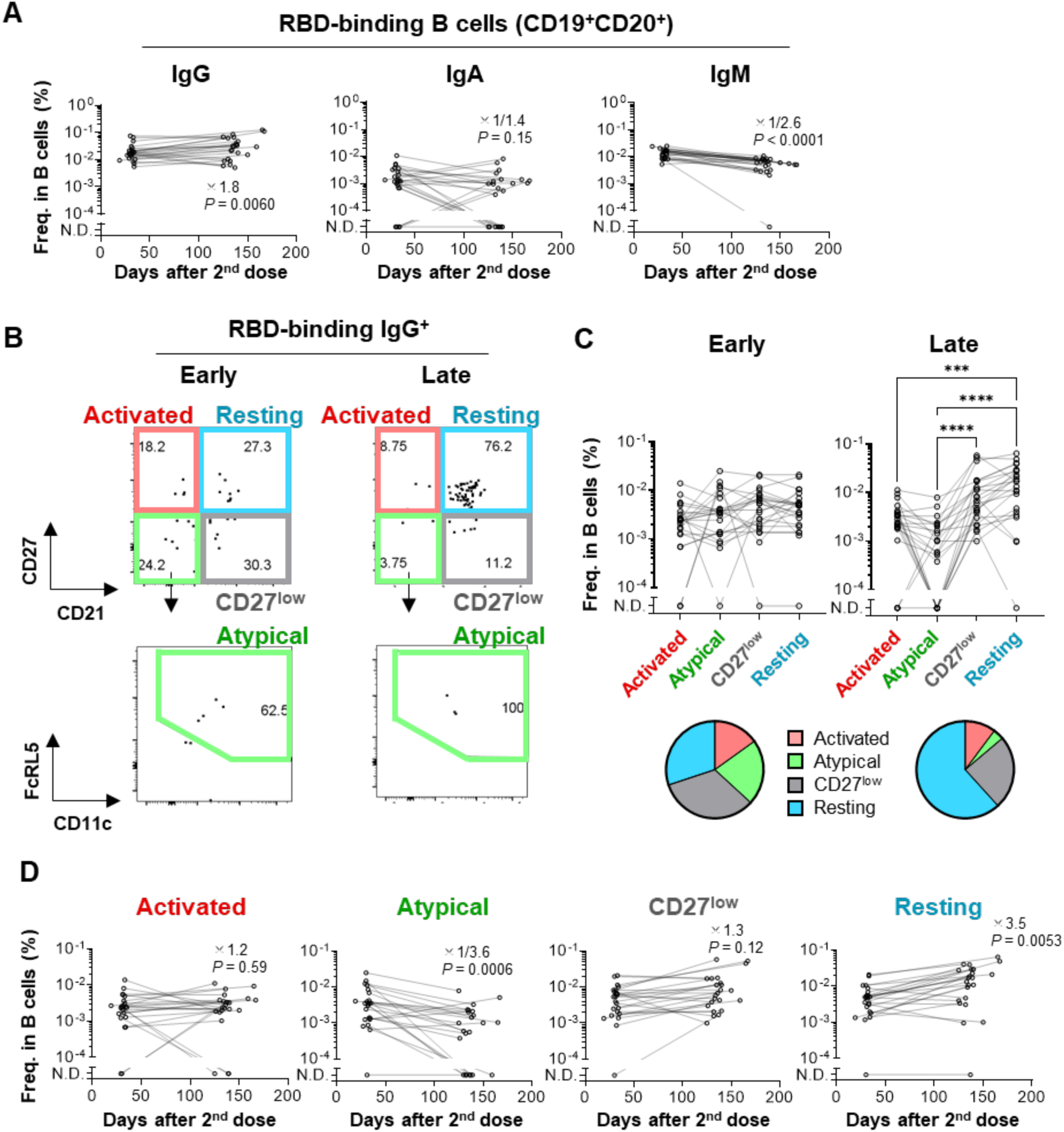
Memory B cell dynamics after mRNA vaccination. (**A**) Longitudinal changes in the number of Wuhan RBD-binding B cells (spike/RBD-binding CD19^+^CD20^+^ cells) expressing IgG, IgA, or IgM were enumerated using flow cytometry. (**B**) RBD-binding IgG^+^ B cells (spike/RBD-binding CD19^+^CD20^+^IgG^+^ cells) were subdivided into four B_mem_ subsets, including activated (CD21^-^CD27^high^), atypical (CD21^-^ CD27^low^CD11c^+^FcRL5^+^), CD27^low^ (CD21^+^CD27l^low^), and resting (CD21^+^CD27^high^) B_mem_ cells. (**C**) The number of the four B_mem_ subsets were determined as gated in (**B**). In the pie charts, frequency of B_mem_ subsets among RBD-binding IgG^+^ B cells were plotted. (**D**) Longitudinal changes in the numbers of RBD-binding IgG^+^ B_mem_ subsets were analyzed with flow cytometry. Median of the fold-changes with *P* values analyzed with the Wilcoxon test (IgG and IgM in A) or fold-change of median with *P* values analyzed with the Mann-Whitney (IgA in A and all the subsets in D) are indicated. For the changing rates, statistical analysis was performed with the Kruskal-Wallis test (\*\*\**P* < 0.001, \*\*\*\**P* < 0.0001).

B_mem_ cell dynamics following mRNA vaccination were tracked in more details. In this study, four subsets of RBD-binding IgG^+^ B_mem_ cells were separately enumerated based on surface markers (Fig. 3B). At 1 month after vaccination (early), four subsets were comparably induced in the vaccinees (Fig. 3C). However, up to 4.9 months after vaccination, the number of resting B_mem_ subset increased by 3.5-fold, whereas the atypical B_mem_ subset decreased in number during the same time period (Fig. 3D), leading to the expanded composition of the resting B_mem_ subset among the IgG^+^ B_mem_ compartment in place of the reduction of the atypical B_mem_ subset. Thus, we revealed the temporal dynamics of vaccine-elicited B_mem_ subset composition, resulting in the dominance of the resting B_mem_ subset in place of the loss of the atypical B_mem_ subset.

### Cross-reactivity of persistent B_mem_ subset against the variants

B_mem_ cells and plasma cells are clonally selected by the differential affinity threshold; therefore, B_mem_ cells tend to express antibodies with lower affinity, and presumably broader reactivity against the primed antigens than plasma cells (*30–33*). Indeed, B_mem_ cells are more crossreactive against the virus variants than circulating antibodies from plasma cells in animal models (*34, 35*); however, the clinical relevance of this phenomenon remains unknown. Simultaneous labeling with Beta and Delta RBD probes allowed us to quantify cross-reactive B_mem_ cells against these variants (Fig. 4, A and B). Among the Wuhan RBD-binding IgG^+^ B-cells, the majority were indeed cross-reactive against the Beta (median: 73.3% at the early and 79% at the late time points) and Delta (median: 71.4% at the early and 79.3% at the late time points) RBD variants. Then, the cross-reactivity of the resting B_mem_ subset was evaluated as shown in Fig. 4C. Of note, the crossreactivity against the Beta and Delta RBDs among the resting B_mem_ subset increased from the early to the late time points, showing that the resting B_mem_ subset increased variant-reactivity as well as cellularity over time. Indeed, the temporal increase in the variant-reactivity was achieved by the prominent expansion in the numbers of Beta- (4.7-fold) and Delta- (4.9-fold) binding cells compared to those of Wuhan-specific cells (1.6-fold) (Fig. 4D).

**Fig. 4.**
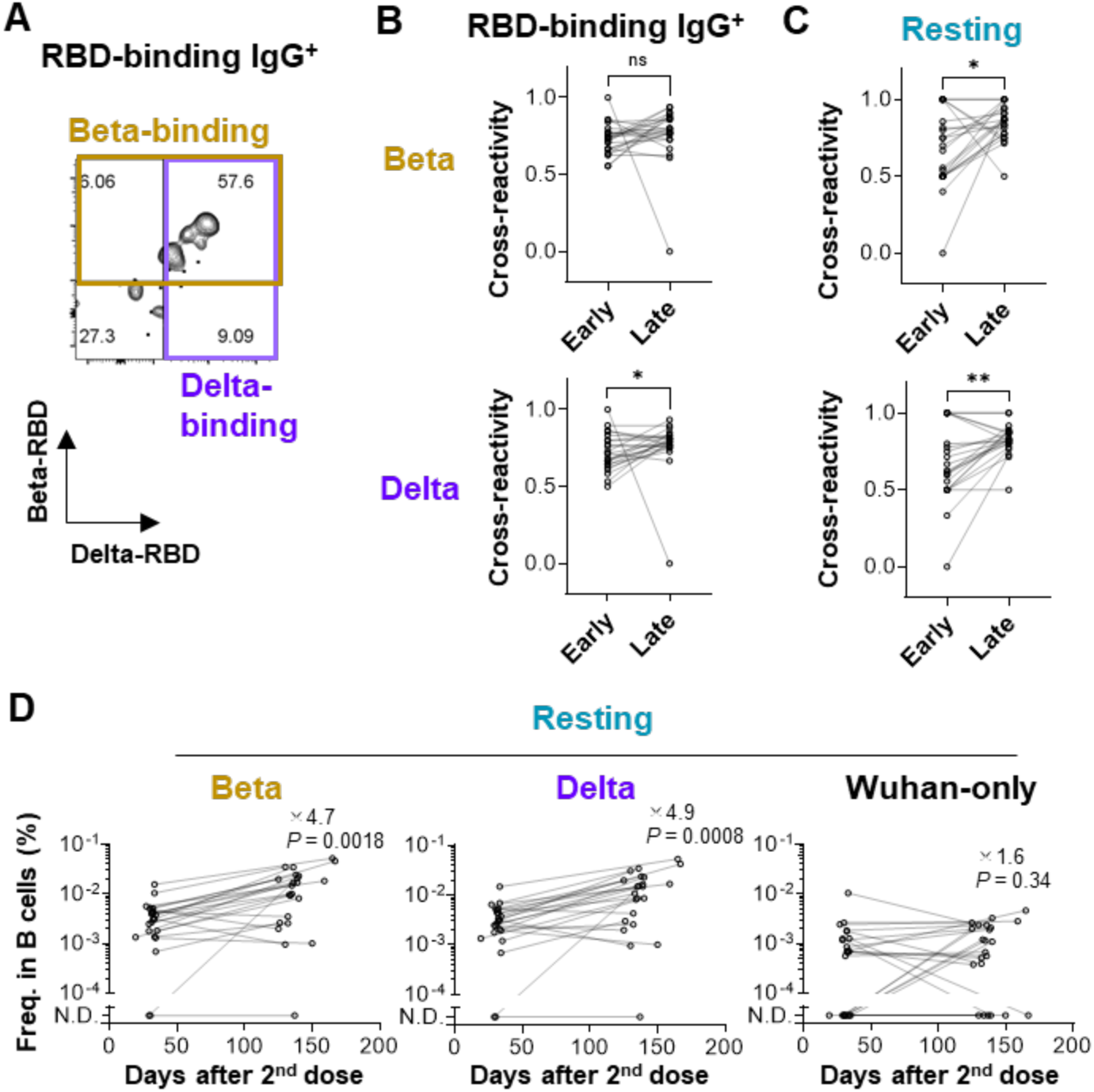
The dynamics of cross-reactive B_mem_ cells with resting phenotype. (**A**) Wuhan RBD-binding IgG^+^ B_mem_ cells were subdivided into Beta- and Delta-binding cells. (**B**) Frequency of the cells that bind the Beta (upper panels) or the Delta (lower panels) RBDs among the Wuhan RBD-binding IgG^+^ B_mem_ cells were plotted. (**C**) Cross-reactivity to the Beta (upper) and Delta (lower) RBD of resting B_mem_ cells was measured. (**D**) Longitudinal changes in the numbers of IgG^+^ resting B_mem_ cells that bind Wuhan/Beta RBD (Beta), Wuhan/Delta RBD (Delta), and Wuhan RBD only (Wuhan-only) were analyzed with flow cytometry. Statistical analyses were performed with the Wilcoxon test (B, C, and D; \**P* < 0.05, \*\**P* < 0.01). In (D), fold-changes of median from the early to the late time points are described with *P* values.

### Omicron-neutralizing activity of plasma antibody

Following the emergence of the Beta and Delta variants, the antibody-escaping variants continuously emerged, including Mu and Omicron variants. Plasma IgG antibodies binding to a panel of RBD proteins form Alpha, Beta, Delta, Mu, and Omicron variants were quantified by ECLIA in parallel to compare the levels of antigenic change (Fig. 5A). In addition, we generated RBD protein that carries multiple mutations conferring resistance to classes 1, 2, and 3 RBD-binding antibodies, known as PMS20 (*36*). At 1 month after the second vaccine dose, plasma IgG antibodies bound to Beta and Mu RBDs at 2.3-fold reduction and PMS20 RBD at 2.5-fold reduction, respectively, confirming the antigenic alteration in these variant RBDs. More strikingly, however, the concentrations of IgG antibodies to Omicron RBD dropped to less than 1/10 of those to Wuhan RBD at the early time point. Further, Omicron RBD-binding IgG antibodies were almost undetectable by 4.9 months (Fig. 5B). These results demonstrate the most profound antigenic change of Omicron variant among those that emerged so far.

**Fig. 5.**
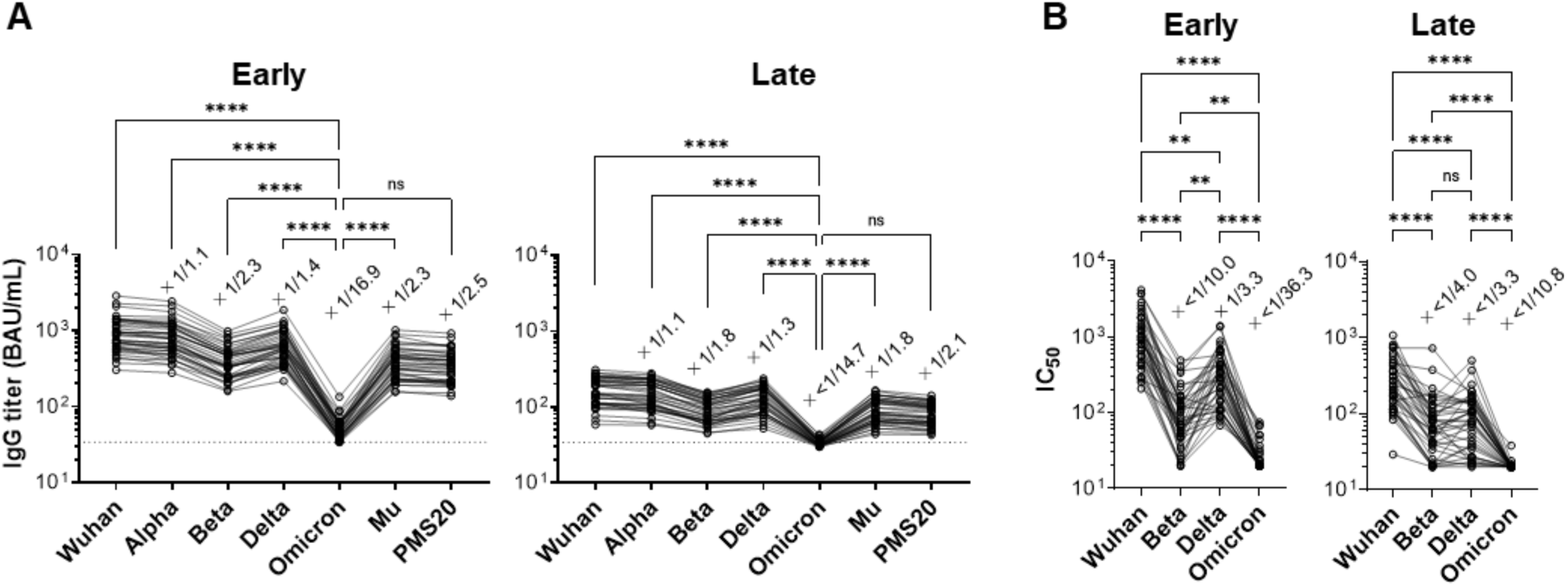
The omicron variant is resistant to plasma neutralizing antibody. (**A**) Plasma IgG titers to mutant RBDs were quantitated with ECLIA. (**B**) Neutralizing antibody titers were quantified using PV expressing Wuhan, Beta, Delta, and Omicron S protein. Median of the fold-changes are indicated on the plots. Statistical analysis was performed with the Friedman test (\*\**P* < 0.01, \*\*\*\**P* < 0.0001, ns, not significant: *P* ≥ 0.05). In (A), only the comparisons with Omicron RBD were indicated.

Next, we examined the resistance of plasma neutralizing antibodies to Omicron variant by PV neutralization assay similar to Fig. 2C. Supporting the recent accumulating evidence (*12–16*) and our binding data (Fig. 5A), the remarkable reduction was observed in the Omicron-neutralizing activity not only in the early time point but also in the late time point (Fig. 5B), showing that antigenic alteration of Omicron variant exceeded the levels which could be adjusted by the temporal maturation of neutralization breadth in plasma antibodies.

### Omicron-neutralizing activities of memory-derived monoclonal antibodies

The preservation of cross-reactive BCRs in the resting B_mem_ subset prompted us to examine the neutralizing activity against the Omicron variant. Flow cytometric analysis allowed us to quantify the percentages of RBD-binding B_mem_ cells that cross-react with the variant RBDs; however, the cross-neutralizing activities could not be assessed through this binding assay. To this end, RBD-binding IgG^+^ B_mem_ cells with resting phenotype were sorted from a total of 23 donors for single-cell culture (Fig. 6A) (*37*). Monoclonal antibody clones secreted into the supernatants were then utilized for screening Wuhan RBD-binding activity by ELISA. Among 98 RBD-binding IgG clones from resting B_mem_ cells, 35 clones were selected as the neutralizing clones through an initial screening (Fig. 6B). Subsequently, 34 clones were confirmed to be the neutralizers by three independent experiments (early, *n* = 9; late, *n* = 25) (Fig. 6C). Thereafter, these IgG clones were subjected to PV-neutralizing assay for assessing the cross-neutralizing activities against the Beta and Omicron variants. Intriguingly, despite the strong resistance of Beta and Omicron variants to plasma neutralizing antibodies, the substantial fraction of memory-derived antibody clones retained the neutralizing activities to the Beta and Omicron variants (Fig. 6C). In total, 21 (59%) and 10 (27%) out of 34 antibodies cross-neutralized Beta and Omicron variants, respectively (Fig. 6D). The distribution of cross-neutralizing antibodies at the early and late time points was roughly comparable (Fig. 6D), although the temporal increase in the neutralization breadth against the Beta and Omicron variants needs to be carefully assessed by larger numbers of antibody panel in the future.

**Fig. 6.**
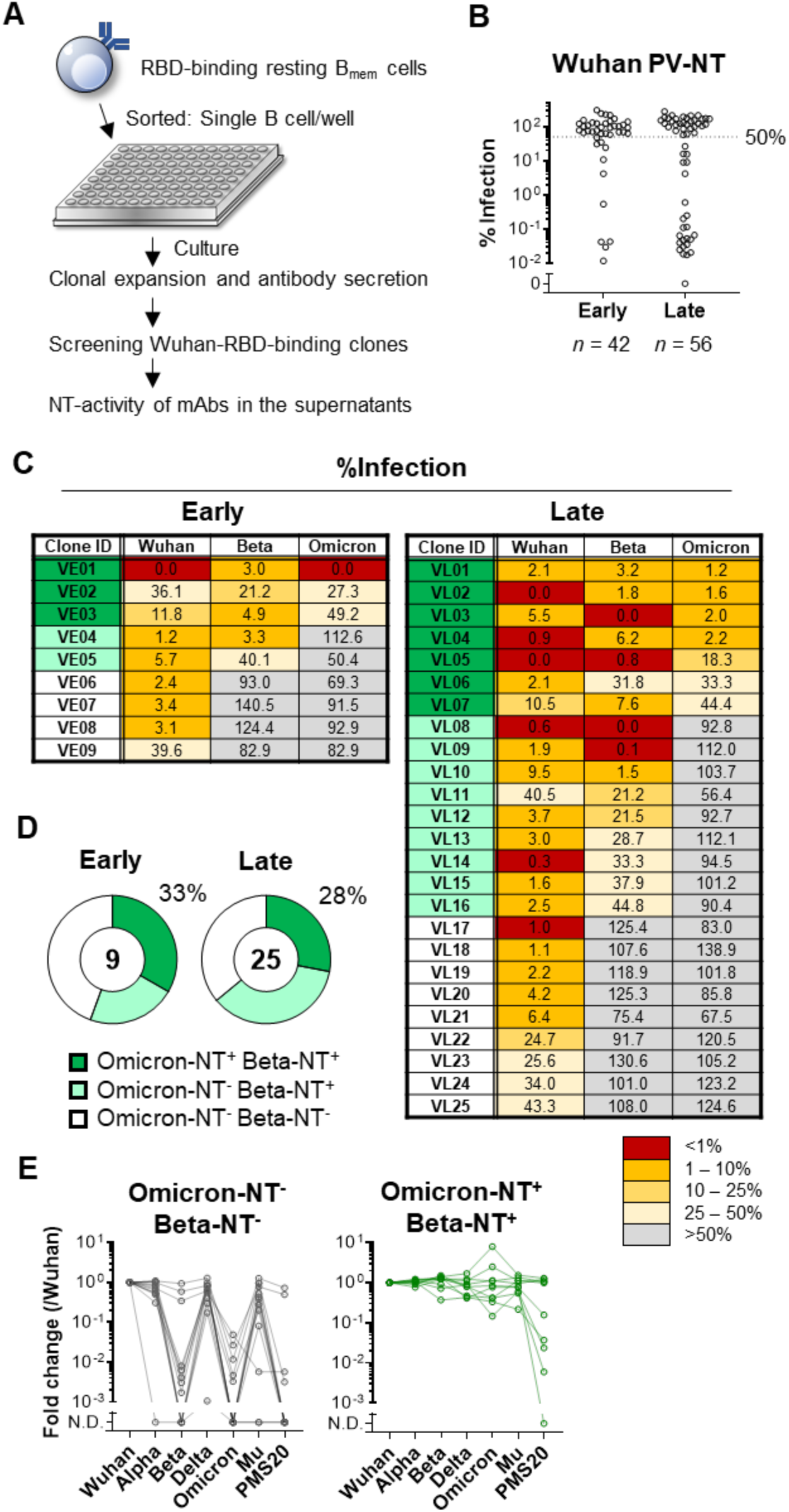
B_mem_ cells preserve Omicron-neutralizing antibody repertoire. (**A**) Schematic diagram of the experimental workflow for assessing the neutralizing (NT) activity of monoclonal antibodies expressed by B_mem_ cells. (**B**) NT activities of individual monoclonal antibody were plotted as % infection calculated as the signals detected by individual antibody out of those without antibody. The antibodies with less than 50% of values were considered as neutralizing antibodies. (**C**) Mean % infection of monoclonal antibody clones with Wuhan-neutralizing activity from early (*n* = 9) and late (*n* = 25) time points of three independent experiments were indicated. (**D**) Pie charts represent the ratios of monoclonal antibody clones with Beta- and Omicron-neutralizing activity. (**E**) Binding breadth of monoclonal antibody clones listed in (C) was quantified with ECLIA. Signals to each RBD were normalized to those of S309 monoclonal antibody and fold changes to Wuhan RBD were calculated. Clones neutralizing Wuhan-only (Omicron-NT^-^ Beta-NT^-^) and Wuhan/Beta/Omicron (Omicron-NT^+^ Beta-NT^+^) were plotted separately.

Finally, the binding breadth of the neutralizing antibody clones to a panel of variant RBDs were assessed by ECLIA similar to Fig. 5A. All of Wuhan-specific neutralizing clones (Omicron^-^ Beta^-^; n = 13) poorly bound to the Omicron RBD but most of them retained the binding to Alpha, Delta, and Mu variants (Fig. 6E). They failed to bind Beta RBD except three clones (VE08, VE09, and VL22) that bound Beta RBD without neutralizing activity (Fig 6, C and E). Two clones (VE08 and VE09) further bound PMS20 RBD, showing the broad RBD binding except Omicron RBD. The partial mismatch between the binding and neutralizing activity is likely, at least partly, due to the better sensitivity in the binding assay than neutralization assay. Nonetheless, the binding of all Omicron^+^Beta^+^ clones (*n* = 10) to Beta and Omicron RBDs clearly confirmed the cross-neutralizing activity. Moreover, half of the Omicron-neutralizing clones showed the strong binding to PMS20 as well. These results highlight the extended breadth of Omicron-neutralizing clones that are durably preserved in the resting B_mem_ subset.

## DISCUSSION

B_mem_ cells have been considered as backups for pre-existing antibodies by plasma cells that provide immediate homologous protection via highly specific, high-affinity antibodies (*6, 38*). Considering the breadth of B_mem_ cells against viral variants, B_mem_ cells have been proposed to potentiate cross-protection that may not be sufficiently afforded by circulating antibodies. However, the clinical relevance of such B_mem_ cell reactivity during protective immunity in humans remains unknown. We demonstrated that the majority of human B_mem_ cells elicited by two doses of mRNA vaccine are able to capture SARS-CoV-2 variants, despite the striking resistance of the variants to pre-existing antibodies (*12–14, 16, 17*). Moreover, such memory cross-reactivity was linked to the cross-neutralizing activities against the variants including highly mutated Omicron variant. These results provide cellular basis behind the profound recall of the variant-neutralizing antibody upon the third dose of the mRNA vaccine (*16, 18, 19*).

mRNA vaccination induces durable B_mem_ cells that express RBD IgG with neutralizing activity. By tracking the B_mem_ cell subsets for 4.9 months after vaccination, we identified the resting subset as the persistent IgG^+^ B-cell subset that dominated among the RBD-binding B cell population over time. Germinal center (GC) responses elicited by mRNA vaccination last more than 100 days with continuous accumulation of somatic hypermutations (*39*). Given the increased numbers of somatic hypermutations in cross-reactive B_mem_ cells (*1*), we speculate that cellular outputs from persistent GC responses are biased to supply cross-reactive B cells that eventually acquire the resting phenotype. The cellular dynamics likely account for the temporal increase in the numbers of the cross-reactive B_mem_ cells with resting phenotype. The neutralization breadth of B_mem_ repertoires to the variants is more expanded after SARS-CoV-2 infection than vaccination (*2*). We speculate that repeated vaccination by mRNA vaccines by itself elicit the cross-neutralizing memory B_mem_ repertoires, albeit at lower levels than infection, but at sufficient levels to recall the broadly neutralizing antibody against the Omicron variant.

Robust recall of the cross-neutralizing antibody by the third dose of vaccine shows the promise to combat against the Omicron variant (*16, 18, 19*). The cross-reactive B_mem_ subset is probably a key cellular component behind the broadened neutralizing antibody responses. The increasing numbers of cross-reactive B_mem_ cells over time support the durability. Moreover, the resting phenotype among the cross-reactive B_mem_ subset is indicative of their ability to produce plasma cells upon restimulation (*40–42*). Cellular and molecular basis underlying the development and function of the cross-reactive B_mem_ subset is important to determine the vaccine strategy that effectively induce and durably maintain the cross-neutralizing antibodies in the vaccinees.

A primary limitation of our study is the lack of longitudinal analysis after the third dose of vaccines in our cohort. Therefore, we cannot determine the extents to which the numbers of cross-neutralizing B_mem_ subset correlate with the magnitudes of cross-neutralizing antibody responses following the third vaccine dose. Owing to the paucity of B_mem_ subsets, detailed phenotypic and transcriptomic characterization of the cross-neutralizing B_mem_ subset is hampered in this study. Finally, we focused on the neutralizing antibodies but non-neutralizing antibodies also confer the protection *in vivo* at least in animal models. The analysis on non-neutralizing and protective antibodies against the variants may generate the distinct outcome owing to the epitope difference.

## MATERIALS AND METHODS

### Human samples

Longitudinal blood samples were collected approximately at days 0, 51, and 161 after the first vaccination with the BNT162b2 mRNA vaccine (Pfizer) from healthcare workers who received two doses of the vaccine at Tokyo Metropolitan Bokutoh Hospital, Japan Community Health Care Organization Tokyo Shinjuku Medical Center, and Showa General Hospital. Blood was collected in Vacutainer CPT tubes (BD Biosciences), followed by centrifugation at 1800 × g for 20 min. Peripheral blood mononuclear cells (PBMCs) were suspended in plasma and harvested into conical tubes, followed by centrifugation at 300 × g for 15 min. The plasma was transferred into another conical tube, whereas the PBMC pellets were washed with PBS three times before cryopreservation in Cell Banker 1 (Zenoaq). The plasma samples were further centrifuged at 800 × g for 15 min and transferred into another tube to completely remove the PBMCs. The plasma samples were heat-inactivated at 56 °C for 30 min before use. Nucleocapsid antibody was analyzed by cobas e411 plus (Roche) with Elecsys Anti-SARS-CoV-2 (Roche), and <1 was evaluated as seronegative according to the manufacturer’s instructions. The present study was approved by the Institutional Review Board of the National Institute of Infectious Diseases (#1321) and was performed according to the Declaration of Helsinki. All volunteers provided written informed consent prior to the enrollment.

### Recombinant RBD preparation

The original and variant RBDs were prepared as previously described (*20*). Briefly, the human codon-optimized DNA sequence encoding amino acids 331-529 of the SARS-CoV-2 spike (GenBank: MN994467) with an N-terminal signal peptide sequence (MIHSVFLLMFLLTPTESYVD) and C-terminal avi-tag and histidine-tag were cloned into the pCAGGS vector. The vectors encoding variant RBDs bearing 501Y mutation (alpha strain), K417N/E484K/N501Y mutations (beta strain), L452R/T478K mutations (delta strain), G339D/S371L/S373P/S375F/K417N/N440K/G446S/S477N/T478K/E484A/Q493R/G496S/Q49 8R/N501Y/Y505H mutations (omicron strain), R346K/E484K/N501Y mutations (mu strain), R346S/K417N/N440K/V445E/L455R/A475V/E484K/M501Y mutations (PMS20) (*36*) were generated in the same frame. The vectors were transfected into Expi293F cells according to the manufacturer’s instructions, and recombinant proteins produced in the supernatant were purified using Ni-NTA agarose (QIAGEN). To biotinylate RBD proteins, the RBD expression vectors were co-transfected into Expi293F cells together with the secreted BirA-Flag plasmid (Addgene) in the presence of 100 μM biotin.

### ECLIA

Plasma titers for SARS-CoV-2 variant RBDs were measured using v-plex and u-plex kits (Meso Scale Discovery) according to the manufacturer’s instructions. Briefly, plates precoated with RBDs (SARS-CoV-2 Panel 11, Meso Scale Discovery) were incubated with MSD Blocker A reagent at room temperature for 1 h at 700 rpm. The plates were washed with PBS supplemented with 0.05% Tween-20 (washing buffer) three times and incubated with samples diluted in MSD Diluent 100 (Meso Scale Discovery) at room temperature for 2 h at 700 rpm. The plates were washed with washing buffer three times, followed by incubation with sulfo-tag-conjugated antihuman IgG or IgA (Meso Scale Discovery) at room temperature for 1 h at 700 rpm, and then the plates were washed three times with the washing buffer. The plates were treated with MSD Gold read buffer B (Meso Scale Discovery) and electrochemiluminescence was immediately determined with MESOQuickPlex SQ 120 (Meso Scale Discovery). Plasma IgG titers were calculated using the reference standard (Meso Scale Discovery) and were converted to international unit (BAU/mL) using the converting unit provided by the manufacturer. To obtain comparable titers of IgA to those of IgG, we normalized the IgA titers using a converting unit, which was calculated from the binding curves of CR3022 monoclonal antibodies prepared as human IgG1 and human IgA1 isotypes (*20, 43*).

For the u-plex assay, we conjugated the biotinylated RBDs to linker proteins of u-plex development pack (Meso Scale Discovery), according to the manufacturer’s instruction. Briefly, the biotinylated proteins were incubated with the linker proteins at room temperature for 30 min followed by incubation with Stop solution at room temperature for 30 min. The linker-conjugated RBDs were mixed and added to u-plex plates. The plates were incubated at 4 °C overnight and then washed with the washing buffer. Thereafter, the same procedure as the v-plex assay was performed for quantifying plasma IgG and monoclonal IgG antibodies.

### Neutralization assay

Neutralization assays were performed as previously described (*20*). For authentic viral neutralization, plasma samples were serially diluted (2-fold dilutions starting from 1:5) in high-glucose DMEM supplemented with 2% FBS and 100 U/mL penicillin/streptomycin and were mixed with 100 TCID50 SARS-CoV-2 viruses, namely WK-521 (hCoV-19/Japan/TY-WK-521/2020, Wuhan strain), TY8-612 (hCoV-19/Japan/TY8-612/2021, Beta variant), and TY11-927 (hCoV-19/Japan/TY11-927-P1/2021, Delta variant), followed by incubation at 37°C for 1 h. The virus-plasma mixtures were placed on VeroE6/TMPRSS2 cells (JCRB1819) seeded in 96-well plates and cultured at 37 °C with 5% CO_2_ for 5 days. After the culture, the cells were fixed with 20% formalin (Fujifilm Wako Pure Chemicals) and were stained with crystal violet solution (Sigma-Aldrich). The mean cut-off dilution index with > 50% cytopathic effect from 2 to 4 multiplicate series was presented as the neutralizing titer.

For the pseudovirus-neutralization assay, VSV-pseudoviruses bearing SARS-CoV-2 spike protein were generated as previously described (*44*). Briefly, cDNAs encoding the spike proteins of SARS-CoV-2 viruses, including WK-521 (Wuhan strain), TY8-612 (Beta variant), TY11-927 (Delta variant), and TY38-873 (hCoV-19/Japan/TY38-873P0/2021, Omicron variant) were cloned into the pCAGGS expression vector with a 19 aa truncation at the C-terminus of the spike. The plasmid vectors were transfected into 293T cells followed by infection with 0.5 MOI of G-complemented VSVΔG/Luc 24 h after the transfection (*45*), and then the uninfected viruses were washed out. After 24 h of culture, the supernatants with the VSV-pseudoviruses were harvested, centrifuged to remove cell debris, and stored at −80 °C until conducting the neutralization assay. For the assay, the pseudoviruses were mixed with an equal volume of serially diluted plasma (5-fold dilutions starting from 1:10 dilution) or B cell culture supernatants diluted at 1:4 ratio, and were incubated for 1 h at 37 °C. The mixtures were inoculated on VeroE6/TMPRSS2 cells seeded in 96-well solid white flat-bottom plates (Corning) and incubated at 37 °C for 24 h in a humid atmosphere containing 5% CO_2_. Luciferase activity in the cells was measured using the Bright-Glo luciferase assay system (Promega) using a GloMax Navigator Microplate Luminometer (Promega). Half-maximal inhibitory concentration (IC50) were calculated using Prism 9 (GraphPad).

### ELISA

ELISA was performed as previously described (*20*). Briefly, F96 Maxisorp Nunc-ELISA plates (Thermo Fisher Scientific) were incubated with 2 μg/mL of the recombinant RBD or antihuman IgG Fab at 4 °C overnight. The plates were washed with PBS containing 0.05% Tween-20 and blocked with PBS supplemented with 1% BSA. After the blocking buffer was discarded, samples and standards were placed into the wells, and the plates were incubated for 2 h at room temperature or overnight at 4 °C. After washing the plates, HRP-conjugated goat anti-human IgG (Southern Biotech) was added to the wells, and the plates were incubated for 1 h at room temperature After washing, the OPD substrate (Sigma) was added to the wells followed by stopping the reaction with 2N H_2_SO_4_, and the absorbance at 490 nm was determined with an Epoch2 microplate reader (Biotek).

### Flow cytometry

Avi-tag-biotinylated spike and RBD were incubated with subsequent fluorochrome-labeled streptavidin at 4:1.5 ratio overnight at 4 °C as follows: spike with APC-streptavidin (Invitrogen), Wuhan RBD with PerCP-Cy5.5-streptavidin (BioLegend), Beta-RBD with BUV661-streptavidin (BD Biosciences), and Delta RBD with PE-streptavidin (Invitrogen). PBMCs were thawed at 37 °C and immediately washed with DMEM supplemented with 2% FBS, followed by staining with the spike/RBD probes in DMEM supplemented with 2% FBS and 10 μM biotin for 30 min at room temperature. The cells were washed with the medium and stained with subsequent antibodies/reagents using the Brilliant Stain Buffer Plus (BD Biosciences) for 30 min at room temperature: FITC-conjugated anti-IgA (polyclonal rabbit F(ab’)2, Dako), BV421-conjugated anti-IgG (G18-145, BD Biosciences), BV510-conjugated anti-human CD2 (RPA-2.10, BioLegend), BV510-conjugated anti-human CD4 (RPA-T4, BioLegend), BV510-conjugated anti-human CD10 (HI10a, BioLegend), BV510-conjugated anti human CD14 (M5E2, BioLegend), LIVE/DEAD Fixable Yellow Dead Cell Stain kit (Thermo Fisher Scientific), BV605-conjugated anti CD27 (O323, BD Biosciences), BV650-conjugated anti-FcRL5 (509F6, BD Biosciences), BUV395-conjugated anti-CD19 (HIB19, BD Biosciences), BUV496-conjugated anti-CD20 (2H7, BD Biosciences), BUV563-conjugated anti-IgM (UCH-B1, BD Biosciences), BUV615-conjugated anti-CD11c (3.9, BD Biosciences), and BUV737-conjugated anti-CD21 (B-ly4, BD Biosciences). The cells were washed with DMEM supplemented with 2% FBS, followed by resuspension in the medium and flow cytometry with FACS Symphony S6 (BD Biosciences).

### Single B cell culture

A single B cell culture, called Nojima-culture, was performed (*37*). Single RBD-reactive resting B_mem_ cells were sorted into 96-well plates containing RPMI1640 medium supplemented with 10% FBS, 55 μM 2-ME, 100 Units/mL penicillin, 100 μg/mL streptomycin, 10 mM HEPES, 1 mM sodium pyruvate, 1% MEM NEAA, 50 ng/mL recombinant human IL-2 (Peprotech), 10 ng/mL recombinant human IL-4 (Peprotech), 10 ng/mL recombinant human IL-21 (Peprotech), 10 ng/mL recombinant human BAFF (Peprotech), and pre-cultured MS40L-low feeder cells at one cell per well using FACS Symphony S6 (BD Biosciences). The plates were incubated at 37 °C in a humid atmosphere with 5% CO_2_. The medium was half-replenished on days 4, 8, 12, 15, and 21, and the supernatants were harvested on day 24.

### Statistical analysis

The numerical data were statistically analyzed and visualized with GraphPad Prism 9 software (GraphPad). Flow cytometry data were analyzed using FlowJo software (BD Biosciences).

## Data and Code Availability

All data needed to support the conclusion of this manuscript are included in the main text and Supplementary materials.

## Acknowledgments

We thank Akira Dosaka, Eriko Izumiyama, Rieko Iwaki, Megumi Koda, Masataka Tokita, Kazuma Ariga, Naoka Yoshida, Kazuko Isoyama, and Ryoko Itami at NIID for their technical support. This work was supported by Japan Agency for Medical Research and Development grant JP19fk0108104 (YT), JP20fk0108104 (YT), JP20fk0108534 (YT), JP21fk0108534 (YT).

## Author contributions

Conceptualization: RK, YA, SM, TO, YT

Methodology: RK, YA, SM, TO, SF

Investigation: RK, YA, SM, TO, TN, KT, KT, LS, TT, AN, MS, KO, FN, HS, TM, MI

Funding acquisition: YT

Project administration: YT

Supervision: YT

Writing – original draft: RK, YA, SM, TO, SF, YT

Writing – review & editing: RK, SM, YT

